# DNA methylation directs nucleosome positioning in RNA-mediated transcriptional silencing

**DOI:** 10.1101/2020.10.29.359794

**Authors:** M. Hafiz Rothi, Shriya Sethuraman, Jakub Dolata, Alan P Boyle, Andrzej T Wierzbicki

## Abstract

Repressive chromatin modifications are instrumental in regulation of gene expression and transposon silencing. In *Arabidopsis thaliana*, transcriptional silencing is performed by the RNA-directed DNA methylation (RdDM) pathway. In this process, two specialized RNA polymerases, Pol IV and Pol V, produce non-coding RNAs, which recruit several RNA-binding proteins and lead to the establishment of repressive chromatin marks. An important feature of chromatin is nucleosome positioning, which has also been implicated in RdDM. We show that RdDM affects nucleosomes via the SWI/SNF chromatin remodeling complex. This leads to the establishment of nucleosomes on methylated regions, which counteracts the general depletion of DNA methylation on nucleosomal regions. Nucleosome placement by RdDM has no detectable effects on the pattern of DNA methylation. Instead, DNA methylation by RdDM and other pathways affects nucleosome positioning. We propose a model where DNA methylation serves as one of the determinants of nucleosome positioning.

## INTRODUCTION

Transcriptional gene silencing (TGS) pathways play an important role in maintaining genomic integrity in eukaryotes. This is achieved through repressive chromatin modifications, which are specifically targeted to silence transposable elements (TE) present in the genome. TGS pathways are conserved in fungi, animal and plant kingdoms, denoting their importance in the control of genome stability (Du et al., 2015). In plants, TGS is established and partially maintained through RNA-directed DNA methylation (RdDM), which consists of two major steps, biogenesis of short interfering RNA (siRNA) and *de novo* DNA methylation (Matzke and Mosher, 2014).

In the first step, RNA polymerase IV (Pol IV) binds to loci targeted for silencing and produces noncoding RNA, which is then converted into a double-stranded form (dsRNA) by RNA-dependent RNA polymerase 2 (RDR2) and cleaved into 24-nucleotide siRNA by DICER-LIKE 3 (DCL3) (Blevins et al., 2015; Law et al., 2013; Mette et al., 1999, 2000; Zhai et al., 2015). siRNAs are then incorporated into ARGONAUTE 4 (AGO4) and other related AGOs, forming AGO-siRNA complexes (Ye et al., 2012). In the second step, RNA polymerase V (Pol V) produces long noncoding RNA (lncRNA) that acts as a scaffold or otherwise helps recruit downstream effectors (Lahmy et al., 2016; Wierzbicki et al., 2008, 2009). The AGO4-siRNA complex is recruited to Pol V-transcribed loci leading to stepwise binding of INVOLVED IN *DE NOVO* 2 (IDN2) and DOMAINS REARRANGED METHYLTRANSFERASE 2 (DRM2) which deposits DNA methylation (Ausin et al., 2009; Böhmdorfer et al., 2014; Cao and Jacobsen, 2002; Wierzbicki et al., 2009; Zilberman et al., 2003). However, the mechanisms by which DNA methylation and other repressive features of chromatin contribute to transcriptional gene silencing are not fully understood.

RNA-directed DNA methylation is functionally intertwined with nucleosome modifications and positioning. This includes the involvement of pre-existing histone modification and putative chromatin remodelers in recruitment of both Pol IV and Pol V (Johnson et al., 2014; Law et al., 2013), as well as the establishment of repressive histone modifications and nucleosome positioning in the second step of RdDM (Liu et al., 2016; Stroud et al., 2014; Wierzbicki et al., 2008; Yang et al., 2020a; Zhu et al., 2013). The involvement of active chromatin remodeling in transcriptional silencing by RdDM was suggested by an interaction of IDN2 with SWITCH 3B (SWI3B), a subunit of the Switch/Sucrose Non Fermenting (SWI/SNF) chromatin remodeling complex (Zhu et al., 2013). Subunits of this complex also interact with other silencing factors, including HISTONE DEACETYLASE 6 (HDA6) and MICRORCHIDIA 6 (MORC6), which indicates that SWI/SNF may be involved in various aspects of gene silencing (Liu et al., 2016; Yang et al., 2020a). This is consistent with this complex being multifunctional and affecting not only gene silencing but also various other aspects of plant gene regulation (Archacki et al., 2017; Brzeski et al., 1999; Han et al., 2012; Li et al., 2016; Sacharowski et al., 2015; Sarnowski et al., 2005; Wagner and Meyerowitz, 2002).

There are several indications that nucleosome positioning and DNA methylation are somehow connected throughout plant genomes (Chodavarapu et al., 2010; Huff and Zilberman, 2014; Lyons and Zilberman, 2017). However, the exact nature of this connection varies depending on species and genomic regions tested (Huff and Zilberman, 2014; Lyons and Zilberman, 2017). In Arabidopsis, nucleosomes determined by MNase digestion protections have been reported to generally correlate with DNA methylation (Chodavarapu et al., 2010). However, the opposite correlation exists on a subset of Arabidopsis nucleosomes and throughout genomes of certain other species (Huff and Zilberman, 2014; Lyons and Zilberman, 2017). This difference may be explained by the DNA binding of linker histones, which prevent methylation of linker DNA, and by the activity of DDM1, which facilitates methylation of nucleosomal DNA (Lyons and Zilberman, 2017; Zemach et al., 2013). In *Arabidopsis* these two proteins counteract the general preference to methylate linker DNA (Lyons and Zilberman, 2017).

The involvement of linker histones, DDM1 and SWI/SNF in determining the pattern of DNA methylation indicates that the observed connection between nucleosomes and DNA methylation is primarily determined by nucleosomes being inaccessible to DNA methyltransferases. This is supported by *in vitro* data indicating preferential methylation of linker DNA (Felle et al., 2011). However, the opposite relationship has been observed on a few individual loci, where nucleosomes were affected by the *drm2* mutation (Zhu et al., 2013). This indicates that DNA methylation may affect nucleosome positioning. This alternative causality is also supported by some *in vitro* data (Collings et al., 2013). Therefore, the relationship between nucleosomes and DNA methylation remains only partially resolved.

Here, we explore the mechanism by which RdDM affects nucleosome positioning in *Arabidopsis thaliana*. We demonstrate that Pol V and more broadly RdDM affects nucleosomes through the SWI/SNF complex. The SWI/SNF complex is not required for DNA methylation on positioned nucleosomes. Instead, DNA methylation is needed for nucleosome positioning on differentially methylated regions. We propose a model where the RdDM pathway directs nucleosome positioning through DNA methylation to establish transcriptional gene silencing.

## RESULTS

### Pol V affects nucleosomes by a combination of direct and indirect mechanisms

Pol V has been previously shown to affect protection to MNase digestion of certain genomic regions (Zhu et al., 2013). To conclusively attribute these protections to nucleosome positioning, we expanded this experiment by including immunoprecipitation with an anti-H3 antibody (MNase H3 ChIP-seq) in two biological replicates of Col-0 wildtype and *nrpe1*, a mutant of the largest subunit of Pol V [Figure 1A]. We identified 690 nucleosomes stabilized by Pol V, where signal was at least 2-fold higher in Col-0 compared to *nrpe1* with a false discovery rate (FDR) of less than 0.05 [Figure 1B]. We also identified 3082 Pol V destabilized nucleosomes, where signal was at least 2-fold higher in *nrpe1* compared to Col-0 with an FDR of less than 0.05 [Figure S1A]. We validated a subset of Pol V stabilized nucleosomes by locus-specific MNase H3 ChIP-qPCR where we detected a significant decrease in nucleosome signal in *nrpe1* compared to Col-0 wildtype at several tested loci [Figure 1E]. HSP70 was used as a negative control (Kumar and Wigge, 2010).

**Figure 1:**
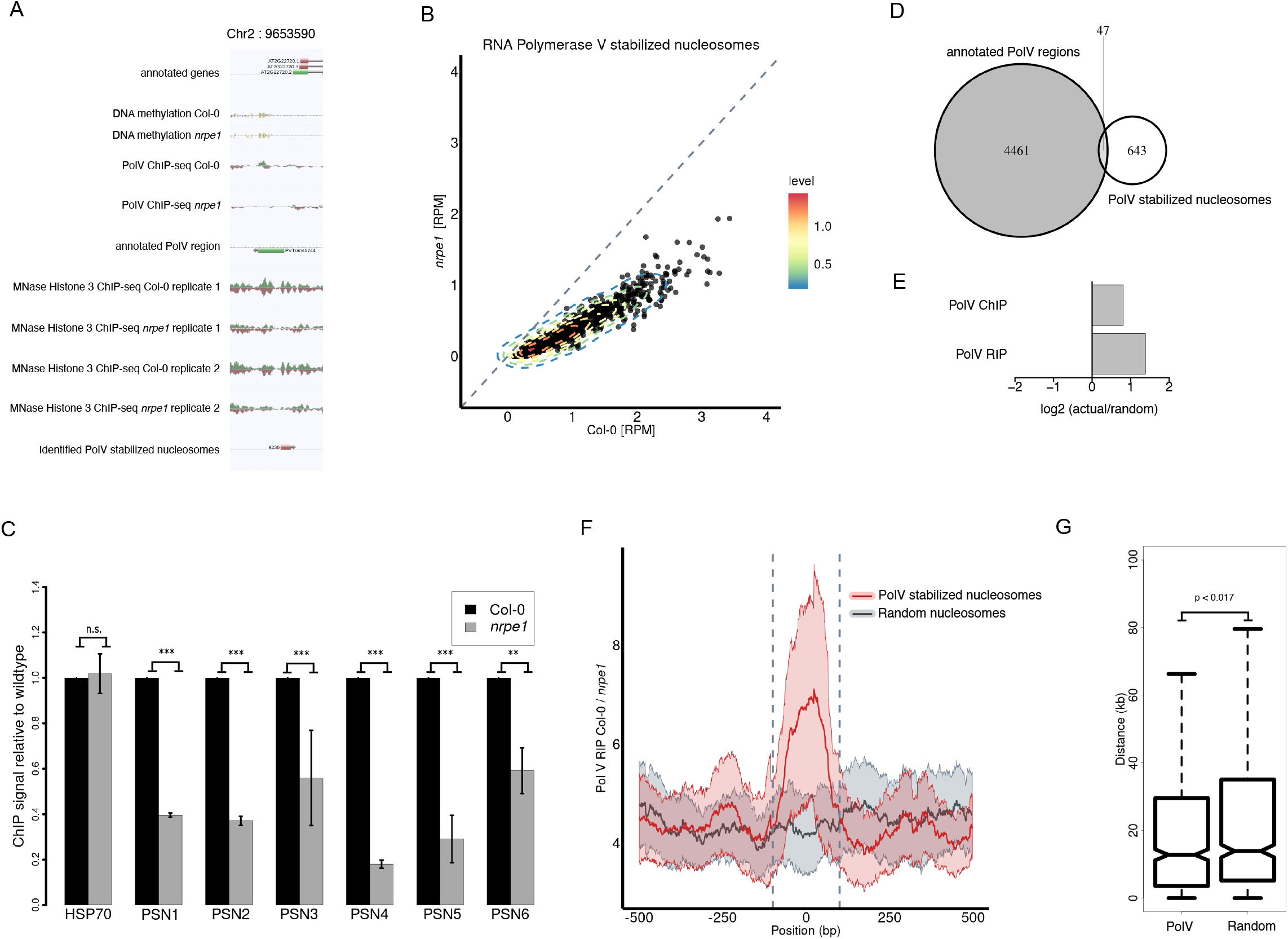
Pol V affects nucleosomes by a combination of direct and indirect mechanisms. A. Genome browser screenshot showing a Pol V stabilized nucleosome. B. Comparison of MNase H3 ChIP-seq signal in Col-0 and *nrpe1* on Pol V stabilized nucleosomes. C. Locus-specific validation of Pol V stabilized nucleosomes. Significance tested using t-test (n.s. = not significant, ** = p-value < 0.01,*** = p-value < 0.001). ChIP signal values were normalized to *ACTIN2* and Col-0 wild-type. Error bars show standard deviations from three biological replicates. D. Overlap between Pol V stabilized nucleosomes and annotated Pol V transcribed regions. E. Enrichment of Pol V stabilized nucleosomes on annotated Pol V transcribed or bound regions (random permutation test; 1000 iterations; p-value < 0.001). F. Pol V RNA immunoprecipitation signal on Pol V stabilized nucleosomes and random nucleosomes. The nucleosomal regions are indicated with vertical dashed lines. G. Distance of Pol V stabilized nucleosomes or random nucleosomes to annotated Pol V transcribed regions.

To test if Pol V stabilized and destabilized nucleosomes are located within Pol V-transcribed regions, we overlapped identified nucleosomes with previously published Pol V-transcribed regions (Böhmdorfer et al., 2016). Pol V stabilized nucleosomes showed a small overlap with annotated Pol V-transcribed regions [Figure 1C], which was still significantly more than expected by chance [Figure 1D]. Consistently, the average level of Pol V transcription on Pol V stabilized nucleosomes was strongly enriched compared to adjacent regions or random sequences [Figure 1F]. Furthermore, like Pol V transcription, Pol V stabilized nucleosomes are enriched in intergenic and promoter regions [Figure S1D-E]. On the other hand, overlaps between Pol V destabilized nucleosomes and annotated Pol V-transcribed regions were less likely than expected by chance [Figure S1B-C]. This indicates that Pol V stabilized nucleosomes are at least partially directly affected by Pol V and its downstream factors while Pol V destabilized nucleosome are most likely affected indirectly.

To determine if Pol V stabilized nucleosomes that do not overlap Pol V-transcribed regions may still be directly affected by Pol V, we measured their distance from Pol V transcribed regions. The average distance between Pol V stabilized nucleosomes and annotated Pol V-transcribed regions was significantly smaller than the average distance between random nucleosomes and annotated Pol V-transcribed regions (Mann-Whitney test, p-value < 0.017) [Figure 1G]. This indicates that Pol V may directly affect nucleosomes that do not overlap annotated Pol V transcripts. Altogether, we conclude that Pol V stabilizes a pool of nucleosomes and at least a subset of those nucleosomes is likely to be directly affected by RdDM.

### Downstream RdDM components affect nucleosome positioning

Involvement of Pol V in nucleosome positioning suggests that other components of the RdDM pathway may also be involved in this process. To test this prediction, we performed MNase-H3 ChIP followed by qPCR in Col-0 wildtype, *nrpe1, ago4-1* and *idn2-1* mutants. We detected a substantial decrease of the nucleosome signals in all three tested mutants compared to wildtype at Pol V stabilized nucleosomes [Figure 2A-G]. While *nrpe1*, as expected, affected all tested nucleosomes, *ago4* and *idn2* had more locus-specific effects [Figure 2A-G]. This indicates that AGO4 and IDN2 both contribute to Pol V-mediated nucleosome positioning. This could be interpreted as evidence that events occurring downstream of Pol V transcription are involved in the observed changes in nucleosome positioning.

**Figure 2:**
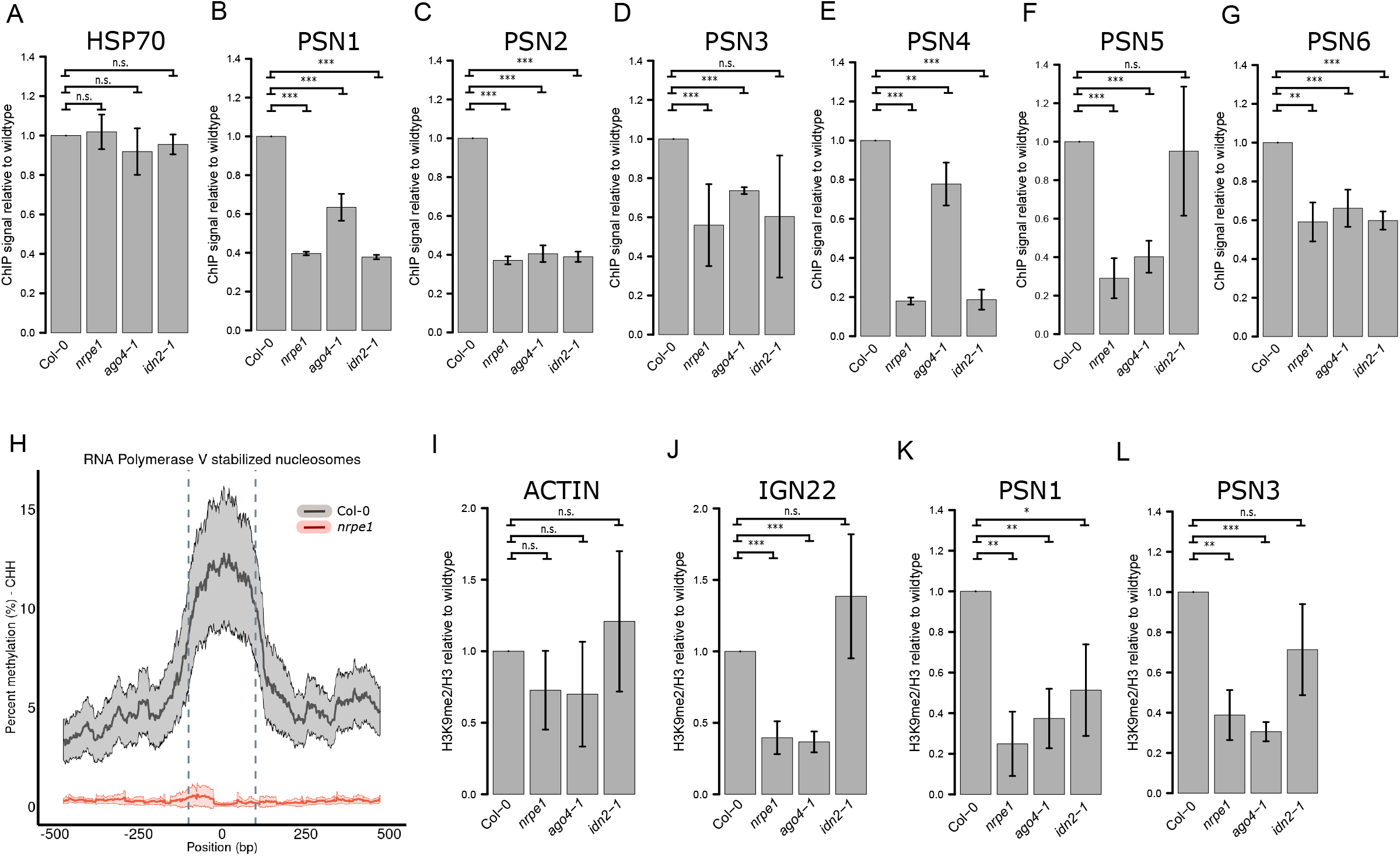
IDN2 connects siRNA and lncRNA to nucleosome positioning. A. – G. Locus-specific analysis of MNase H3-ChIP qPCR levels on Pol V stabilized nucleosomes in Col-0, *nrpe1, ago4-1* and *idn2-1*. Significance tested using t-test (n.s. = not significant, ** = p-value < 0.01,*** = p-value < 0.001). ChIP signal values were normalized to ACTIN2 and Col-0 wild-type. Error bars show standard deviations from three biological replicates. H. Average profile of DNA methylation levels (CHH context) on Pol V stabilized nucleosome dyads. I. – K. Locus-specific analysis of H3K9me2 levels in *ACTIN2, IGN22* and Pol V stabilized nucleosomes in Col-0, *nrpe1, ago4* and *idn2-1*. Significance tested using t-test (n.s. = not significant, * = p-value < 0.05, ** = p-value < 0.01,*** = p-value < 0.001). H3K9me2 ChIP signal values were normalized to H3 and Col-0 wild-type. Error bars show standard deviations from three biological replicates.

### Pol V-stabilized nucleosomes are enriched in repressive chromatin marks

RdDM may affect nucleosome positioning in parallel with establishing repressive chromatin marks like DNA methylation and H3K9 dimethylation (H3K9me2). Alternatively, RdDM may establish nucleosomes and repressive chromatin marks on independent subsets of loci. To distinguish between those possibilities, we performed whole-genome bisulfite sequencing in Col-0 wildtype and *nrpe1* in two biological replicates. We plotted DNA methylation levels in the CHH context at Pol V stabilized nucleosomes and 500 bp adjacent regions [Figure 2H]. CHH DNA methylation was significantly enriched on Pol V stabilized nucleosomes compared to both the adjacent regions and the *nrpe1* mutant [Figure 2H]. To test if this enrichment is also dependent on AGO4 and IDN2, we used previously published whole-genome bisulfite sequencing datasets (Stroud et al., 2013). Likewise, we detected a reduction in the average DNA methylation levels in both *ago4-1* and *idn2-1* [Figure S2A]. These findings indicate that Pol V affects nucleosome positioning in parallel with establishing DNA methylation. This is consistent with genome-wide enrichment of DNA methylation on nucleosomes reported by Chodavarapu *et al*. (2010).

To determine if H3K9me2 is also established in parallel, we performed MNase ChIP-qPCR using anti-H3K9me2 antibody in wildtype, *nrpe1, ago4-1* and *idn2-1* in three biological replicates. We used the anti-H3 antibody as a reference. The levels of H3K9me2 relative to the levels of H3 were significantly reduced on tested Pol V stabilized nucleosomes in *nrpe1* and *ago4* [Figure 2K-L], unchanged on a negative control locus [Figure 2I] and reduced on a positive control locus [Figure 2J]. The *idn2* mutant showed a locus-specific effect, which is consistent with demonstrated partial redundancy of IDN2 and its paralogs (Ausin et al., 2012; Xie et al., 2012). This indicates that at least at the tested loci Pol V affects nucleosome positioning in parallel with establishing H3K9me2. Together, these results indicate that RdDM affects nucleosome positioning in parallel with establishing repressive chromatin marks. This is consistent with a model where Pol V-stabilized nucleosomes are placed and modified by the RdDM pathway.

### Pol V positions nucleosomes via the SWI/SNF chromatin remodeling complex

The presence of a physical interaction between an RdDM factor IDN2 and SWI3B, a subunit of the SWI/SNF chromatin remodeling complex (Zhu et al., 2013), indicates that Pol V may affect nucleosomes via the SWI/SNF complex. To test this possibility, we performed MNase H3 ChIP-seq in Col-0 wildtype and *swi3b* mutant. Although *swi3b* null mutants are embryo lethal (Sarnowski et al., 2005), we took advantage of the well documented observation that *SWI3B* is haploinsufficient (Sarnowski et al., 2002, 2005; Zhu et al., 2013) and used the *swi3b*/+ heterozygous plants. We plotted the average nucleosome signal at Pol V stabilized nucleosomes and adjacent regions [Figure 3A], compared to random nucleosomes identified in Col-0 wild type [Figure 3B]. Nucleosome signal at Pol V stabilized nucleosomes was reduced in *swi3b*/+ [Figure 3AC] but unchanged in adjacent regions [Figure 3AC] and at random nucleosomes [Figure 3BD]. Although the effect observed in *swi3b*/+ was statistically significant, it was much smaller than in *nrpe1* [Figure 3A]. A significant but overall minor reduction in the nucleosomal signal in *swi3b*/+ was confirmed by locus-specific MNase H3 ChIP-qPCR, where we detected small but significant decreases in nucleosome signal in *swi3b*/+ compared to wildtype at tested loci [Figure S3A]. Partial reductions of nucleosome signals in *swi3b*/+ may be explained by the presence of one allele of *SWI3B*, other SWI3 paralogs and other chromatin remodeling complexes. Overall, these results indicate that the SWI/SNF complex only partially contributes to nucleosome positioning by RdDM.

**Figure 3:**
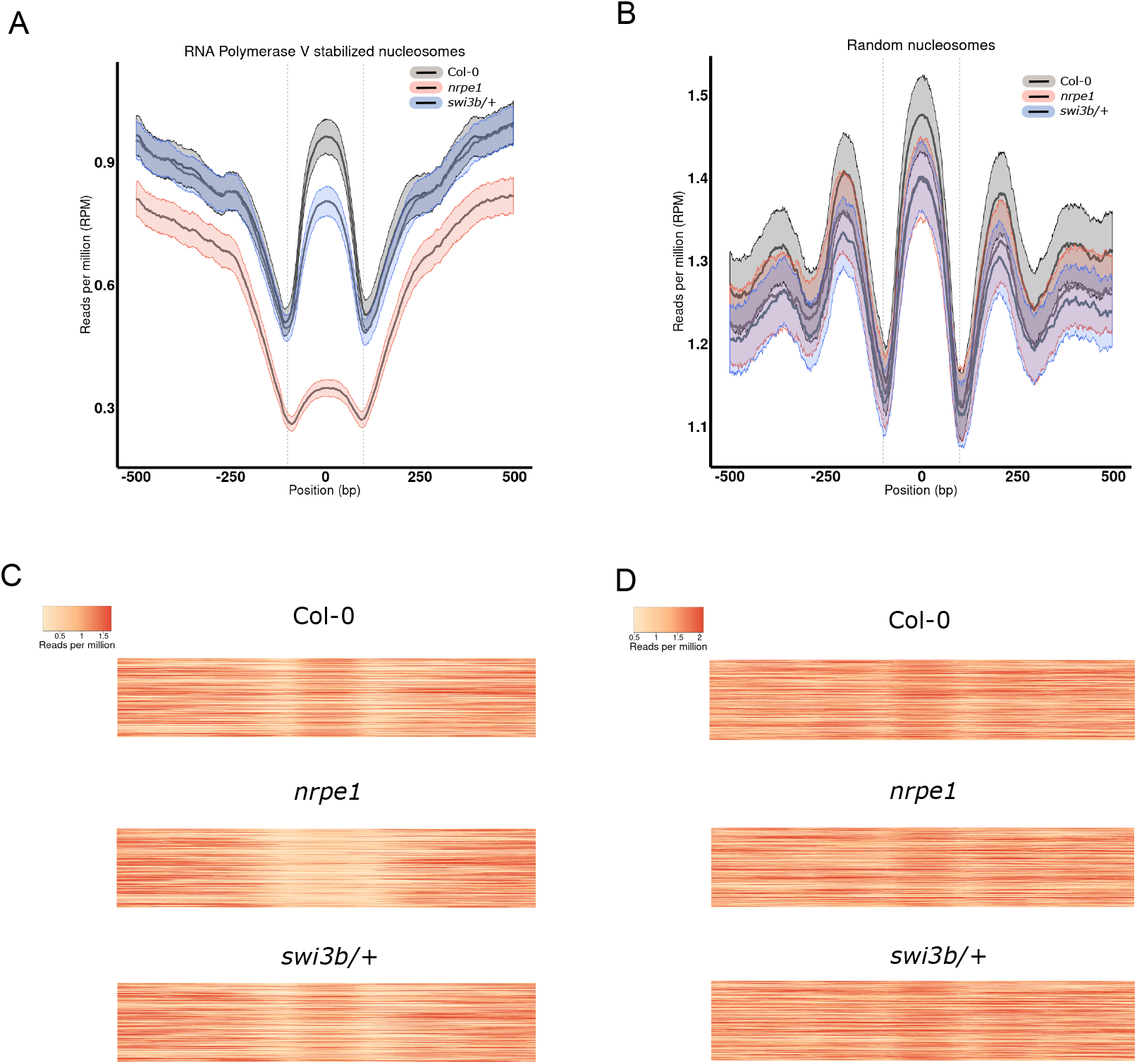
Pol V positions nucleosomes through the SWI/SNF complex. A. Average levels of MNase H3 ChIP-seq signal on Pol V stabilized nucleosomes in Col-0, *nrpe1* and *swi3b*/+. Ribbons indicate confidence intervals with p < 0.05. B. Average levels of MNase H3 ChIP-seq signal on random nucleosomes in Col-0, *nrpe1* and *swi3b*/+. Ribbons indicate confidence intervals with p < 0.05. C. Heatmap of levels of MNase H3 ChIP-seq signal on Pol V stabilized nucleosomes in Col-0, *nrpe1* and *swi3b*/+. D. Heatmap of levels of MNase H3 ChIP-seq signal on random nucleosomes in Col-0, *nrpe1* and *swi3b*/+.

### Preferential methylation of linker DNA

Our observation that RdDM establishes both nucleosome positioning and repressive chromatin marks provides a possible explanation to prior observations that nucleosome positioning and DNA methylation are correlated (Chodavarapu et al., 2010; Lyons and Zilberman, 2017). However, prior studies in Arabidopsis used MNase digestion as the sole basis for the identification of nucleosomes (Chodavarapu et al., 2010; Lyons and Zilberman, 2017) and protections by DNA-binding proteins other than histones remain possible. To eliminate this possibility, we determined nucleosome positioning by MNase H3 ChIP-seq, which relies on MNase protection and binding of histone H3 to DNA to identify nucleosomes. We performed MNase H3 ChIP-seq and whole-genome bisulfite sequencing in two biological replicates of Col-0 wildtype. We first identified all nucleosome positions genome-wide (n=650,610) and measured the average DNA methylation levels in all contexts (CG, CHG and CHH) at nucleosomes and 500 bp adjacent regions. We observed that DNA methylation was enriched on linker regions and depleted on nucleosomes in CHG and CHH sequence contexts [Figure 4B-D]. The CG methylation pattern was more complex, but linkers of neighboring nucleosomes showed strong enrichments in CG context [Figure 4A], which indicates that linkers are preferentially methylated in all sequence contexts. No enrichment was observed on matching random regions [Figure S4A-C]. This indicates that when nucleosomes are identified based on MNase protection and the presence of histone H3, linker regions are enriched in DNA methylation compared to regions occupied by nucleosomes. This correlation is apparent when analyzing all identified nucleosomes [Figure 4A-D] and in most subsets of nucleosomes present on specific genomic regions [Figure S4D-E].

**Figure 4:**
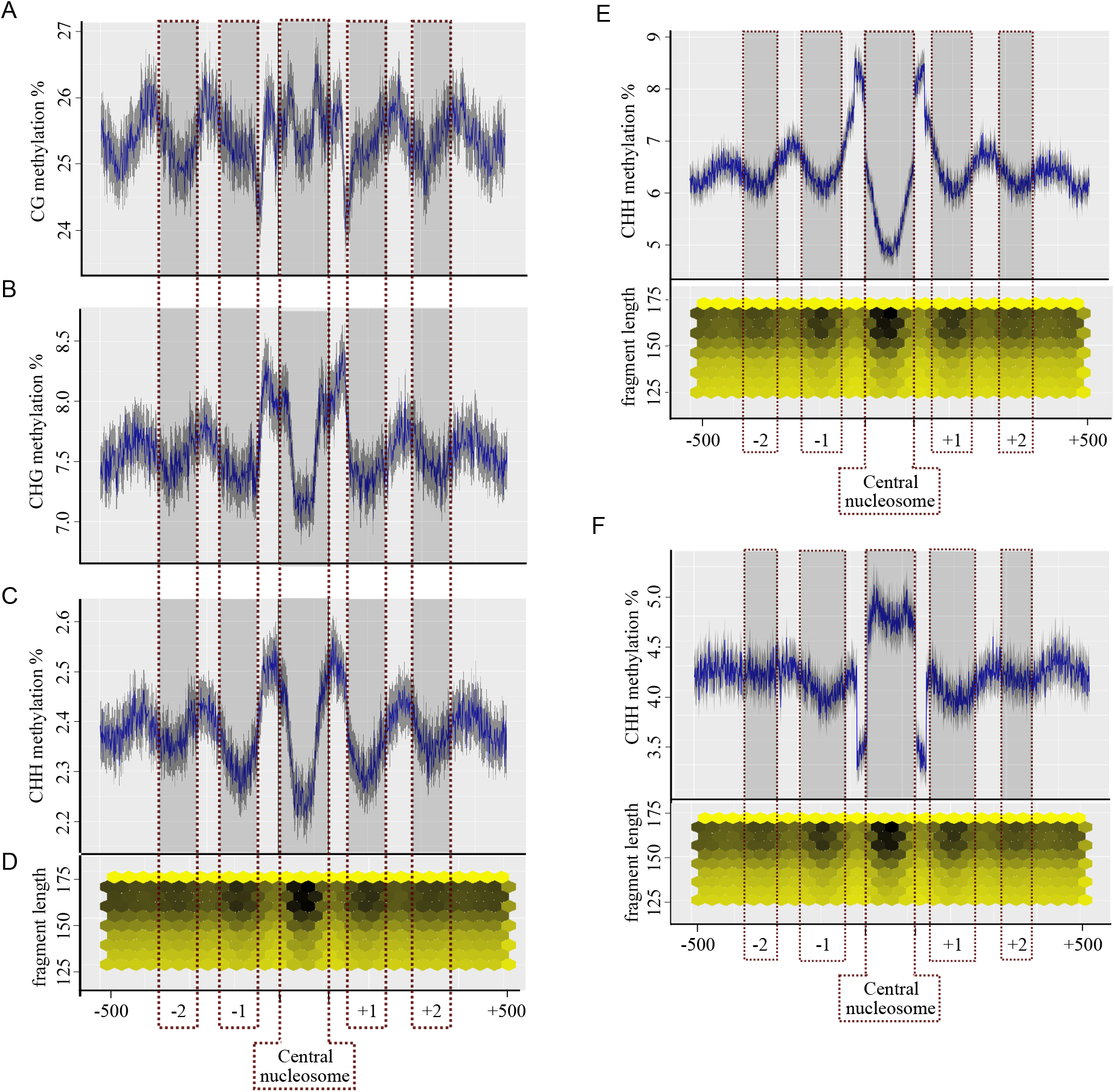
Preferential methylation of linker DNA. A. – C. Average CG (A), CHG (B) and CHH (C) methylation levels on and around all annotated nucleosomes. Dark grey shading indicates the annotated nucleosome and four neighboring nucleosomes. Ribbon indicates confidence intervals with p < 0.05. D. Average MNase H3 ChIP signal levels at and around annotated nucleosomes (X axis) by sequenced fragment length (y axis). E. Average levels of CHH methylation around hypomethylated nucleosomes. Dark grey shading indicates the annotated nucleosome and four neighboring nucleosomes. Ribbon indicates confidence intervals with p < 0.05. Scatterplot below shows average MNase H3 ChIP signal levels at and around hypomethylated nucleosomes (X axis) by sequenced fragment length (y axis). F. Average levels of CHH methylation around hypermethylated nucleosomes. Dark grey shading indicates the annotated nucleosome and four neighboring nucleosomes. Ribbon indicates confidence intervals with p < 0.05. Scatterplot below shows average MNase H3 ChIP signal levels at and around hypermethylated nucleosomes (X axis) by sequenced fragment length (y axis).

Although the average levels of DNA methylation were higher on linker regions than on nucleosomes, a substantial subset of nucleosomes did not follow this general trend, including Pol V stabilized nucleosomes [Figure 2H]. To determine if enrichment of DNA methylation on linkers is generally applicable, we focused on subsets of nucleosomes that show enrichment [Figure 4E] or depletion [Figure 4F] of CHH methylation on their linkers. We then used these subsets to test the enrichment of CHH methylation on linkers of neighboring nucleosomes, which are not expected to be biased by the selection of the central nucleosome. Nucleosomes that follow the general trend, showed the expected enrichment of CHH methylation on linkers of neighboring nucleosomes [Figure 4E]. Interestingly, nucleosomes filtered for depletion of CHH methylation on their linkers, still showed enrichment of CHH methylation on linkers of neighboring nucleosomes [Figure 4F]. This further confirms our observation that on average, DNA methylation is enriched on linker regions. We conclude that while Pol V stabilized nucleosomes are enriched in DNA methylation, the general genome-wide trend is preferential methylation of linker DNA.

### SWI/SNF complex is not required for DNA methylation on positioned nucleosomes

The general trend of methylation depletion on nucleosomal DNA [Figure 4C] is not followed by Pol V stabilized nucleosomes, which are enriched in CHH methylation [Figure 2H]. This indicates that RdDM overrides general preferences of DNA methylation in respect to nucleosome positioning. This may be explained by a hypothesis that nucleosomes positioned by SWI/SNF are preferential substrates for DNA methyltransferases. To test this hypothesis, we assayed DNA methylation by whole-genome bisulfite sequencing in Col-0 wildtype and *swi3b*/+ in two biological replicates. We first analyzed CHH methylation levels on and around Pol V stabilized nucleosomes and observed no change in DNA methylation levels in *swi3b*/+ compared to Col-0 wild type [Figure 5A]. This indicates that the activity of SWI/SNF on Pol V-stabilized nucleosomes has no strong effect on DNA methylation.

**Figure 5:**
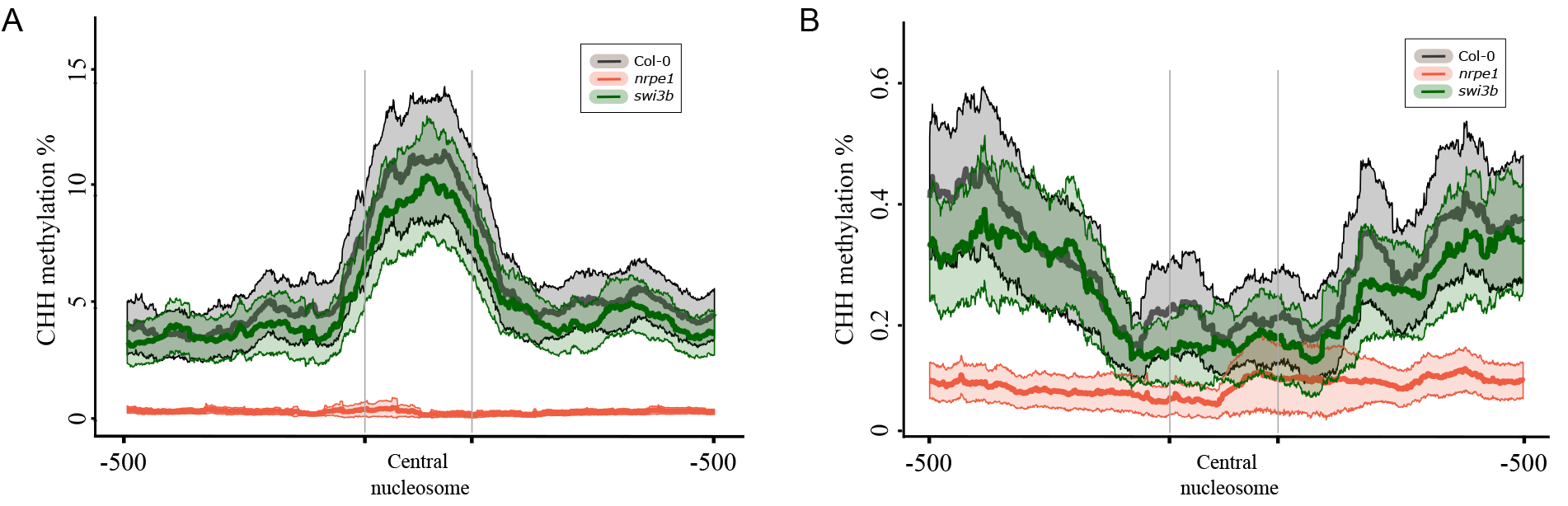
SWI/SNF complex is not required for DNA methylation on positioned nucleosomes. A. Average levels of CHH methylation on and around Pol V stabilized nucleosomes. X axis indicates position (bp). Ribbons indicate confidence intervals with p < 0.05. B. Average levels of CHH methylation on and around SWI3B stabilized nucleosomes. X axis indicates position (bp). Ribbons indicate confidence intervals with p < 0.05.

To further test if nucleosomes positioned by SWI/SNF affect DNA methylation, we identified SWI3B stabilized nucleosomes, which are defined as nucleosomes that have a higher MNase H3 ChIP-seq signal level in wildtype compared to *swi3b*/+ with FDR of less than 0.05. In total, we identified 4089 SWI3B stabilized nucleosomes, where the average nucleosome signal was significantly and reproducibly decreased in *swi3b*/+ [Figure S5A-B]. CHH methylation levels on SWI3B stabilized nucleosomes were not significantly changed in *swi3b*/+ compared to Col-0 wild type [Figure 5B, Figure S5C]. This indicates that changed patterns of nucleosome positioning in *swi3b*/+ do not affect the levels of CHH methylation. This suggests that nucleosomes positioned by the activity of SWI/SNF do not determine the pattern of DNA methylation.

### DNA methylation is needed for positioning nucleosomes at differentially methylated regions

Our observation that correlations between nucleosomes and CHH methylation are not determined by nucleosome positioning suggests an alternative possibility that DNA methylation may participate in determining positions of nucleosomes. To test this prediction, we used previously published datasets (Stroud et al., 2013) to identify differentially methylated regions (DMRs), where CHH methylation is affected by DRM2. We then assayed nucleosome positioning by MNase H3 ChIP-seq in two biological replicates of Col-0 wildtype and *drm2* mutant. At DRM2 DMRs, where lack of DRM2 resulted in strong reductions of CHH methylation [Figure 6A], the nucleosome signal was generally enriched in Col-0 wild type [Figure 6B]. This is consistent with Pol V stabilized nucleosomes being enriched in CHH methylation [Figure 2H]. It is also in agreement with the observation that nucleosomes overlapping DRM2 DMRs behave like Pol V stabilized nucleosomes and are enriched in both CHH methylation and nucleosome signal [Figure S6A-B]. Importantly, in the *drm2* mutant, DRM2 DMRs had a strong reduction in the nucleosome signal [Figure 6B]. This indicates that DNA methylation in the CHH context established by the RdDM pathway affects nucleosome positioning.

**Figure 6:**
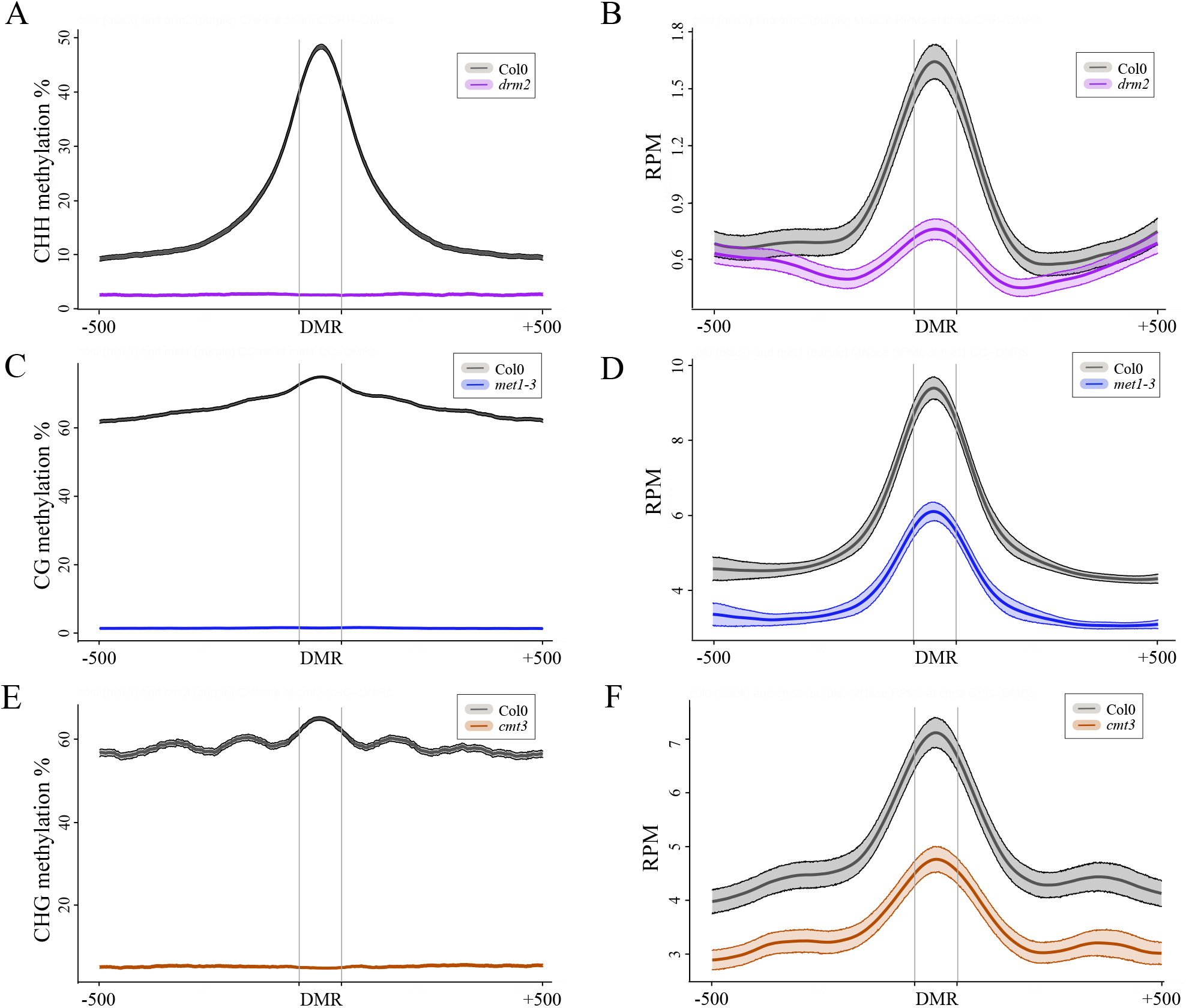
DNA methylation is needed for positioning nucleosomes at differentially methylated regions. A. Average levels of CHH methylation at and around regions that lose CHH methylation in the *drm2* mutant (DRM2 DMRs). Ribbons indicate confidence intervals with p < 0.05. B. Average levels of MNase H3 ChIP signal at and around DRM2 DMRs. Ribbons indicate confidence intervals with p < 0.05. C. Average levels of CG methylation at and around regions that lose CG methylation in the *met1* mutant (MET1 DMRs). Ribbons indicate confidence intervals with p < 0.05. D. Average levels of MNase H3 ChIP signal at and around MET1 DMRs. Ribbons indicate confidence intervals with p < 0.05. E. Average levels of CHG methylation at and around regions that lose CHG methylation in the *cmt3* mutant (CMT3 DMRs). Ribbons indicate confidence intervals with p < 0.05. F. Average levels of MNase H3 ChIP signal at and around CMT3 DMRs. Ribbons indicate confidence intervals with p < 0.05.

To test if this also applies to Pol V stabilized nucleosomes, we plotted the MNase H3 ChIP signal on Pol V stabilized nucleosomes in the *drm2* mutant compared to Col-0 wild type. Consistently with observations on DRM2 DMRs [Figure 6B] and nucleosomes overlapping DRM2 DMRs [Figure S6B], Pol V stabilized nucleosomes also had a reduction in nucleosome signal in the *drm2* mutant [Figure S6C]. This further supports our observation that DNA methylation in the CHH context established by the RdDM pathway affects nucleosome positioning.

To test if DNA methylation in CG and CHG contexts also affects nucleosome positioning we performed similar experiments and analysis in *met1* and *cmt3* mutants. CG and CHG DMRs identified in *met1* and *cmt3*, respectively, were enriched in the nucleosomal signal [Figure 6DF]. At MET1 DMRs, where lack of MET1 resulted in strong reductions of CG methylation [Figure 6C], the nucleosome signal was significantly reduced in *met1* [Figure 6D]. Similarly, at CMT3 DMRs, where lack of CMT3 resulted in strong reductions of CHG methylation [Figure 6E], the nucleosome signal was also significantly reduced in *cmt3* [Figure 6F]. This indicates that DNA methylation affects nucleosome positioning irrespective of the sequence context.

## DISCUSSION

We propose a model, where DNA methylation is a determinant of nucleosome positioning in RdDM. In this model, non-coding transcription by both Pol IV and Pol V leads to the recruitment of AGO4 and IDN2. IDN2 interacts with a subunit of SWI/SNF, which is however not sufficient to affect nucleosome positioning. Instead, the subsequent recruitment of DRM2 and establishment of DNA methylation activates chromatin remodelers and leads to changes in nucleosome positioning. Coordinated establishment of various chromatin marks leads to repression of Pol II promoters within the silenced region of the genome.

The effect of DNA methylation on nucleosome positioning may be explained by distinct intrinsic properties of DNA containing 5-methylcytosines, as suggested by (Collings et al., 2013). Alternatively, DNA methylation may facilitate the recruitment or activation of SWI/SNF, either directly or by the involvement of other proteins that are sensitive to the presence of 5-methylcytosines. Another possibility is that DNA methylation may affect nucleosome positioning by changing the pattern of posttranslational histone modifications. This includes H3K9me2, which may recruit proteins that modulate the activity of chromatin remodelers. This also includes histone deacetylation, which may affect physical properties of the nucleosomes (Yang et al., 2020b; Zhang et al., 2018).

The importance of DNA methylation for nucleosome positioning has a significant impact on our understanding of the RdDM pathway. It argues against the pathway being branched after IDN2 recruitment (Zhu et al., 2013). Instead, it supports the notion that events occurring co-transcriptionally at the sites of Pol V transcription are organized in a stepwise genetic pathway (Böhmdorfer et al., 2014). Although when studied genetically, this pathway appears linear, various steps of the pathway are likely to rely on the cooperative recruitment or activation of subsequent factors. One example of such a connection is the requirement of both IDN2-SWI3B interaction and DNA methylation for nucleosome positioning. Other examples include the recruitment of AGO4, which has been proposed to rely on the interaction of AGO4 with NRPE1 C-terminal domain and with Pol V transcripts (El-Shami et al., 2007; Wierzbicki et al., 2009). Similarly, there is evidence of DRM2 being recruited by interactions with AGO4 and other RdDM factors (Gao et al., 2010; Zhong et al., 2014).

Our model is consistent with the notion that events in the late stages of RdDM lead to a concerted establishment of DNA methylation, posttranslational histone modifications and nucleosome positioning, which together form a repressive chromatin structure. This explains the robustness of transcriptional silencing, where coordinated establishment of various repressive chromatin marks leads to efficient repression of Pol II transcription. It is also consistent with the general difficulty to experimentally tease apart various repressive chromatin modifications established by this pathway.

The involvement of SWI/SNF and nucleosome positioning in RdDM may also be considered in the context of this pathway performing mostly maintenance of silencing. Transcription of heterochromatic regions by Pol IV and Pol V may involve the removal or repositioning of previously positioned nucleosomes. This is supported by the involvement of putative chromatin remodelers in initiation and/or elongation of transcription by both of those polymerases (Kanno et al., 2004; Smith et al., 2007; Wierzbicki et al., 2008; Zhou et al., 2018). Nucleosome positioning established as an outcome of RdDM may serve to re-create the pattern of nucleosomes disrupted by Pol IV and Pol V. *De novo* RdDM in newly inserted TEs is a distinct scenario, since no pre-existing repressive chromatin modifications are expected to exist. The role of nucleosome positioning in this *de novo* process remains unexplored.

The involvement of DNA methylation in determining the pattern of nucleosomes extends beyond RdDM targets. The impact of MET1-dependent CG methylation and CMT3-dependent CHG methylation on stabilizing nucleosomes indicates that DNA methylation may affect nucleosome patterns throughout the genome. This is consistent with findings in other eukaryotes (Collings and Anderson, 2017; Collings et al., 2013). Such a general effect of DNA methylation on nucleosome positioning would counteract the general preference to methylate linkers and contribute to local correlations between nucleosomes and DNA methylation. This property of nucleosomes is consistent with previous reports (Chodavarapu et al., 2010) and may involve the activity of DDM1 (Lyons and Zilberman, 2017). It illustrates the general interdependence between nucleosomes and DNA methylation.

Existing evidence does not support the view that DNA methylation is the primary determinant of the nucleosome pattern. This role remains reserved for a combination of intrinsic factors and active chromatin remodeling. The role of DNA methylation is more limited and probabilistic, clearly visible in meta-analysis of large pools of sequences. Therefore, opposite behaviors of individual loci are expected. Moreover, global losses of DNA methylation in RdDM and DNA methyltransferase mutants may affect the patterns of nucleosomes by a combination of cis- and trans-acting factors, which could only be distinguished using tools targeting DNA methylation to specific loci.

## MATERIALS AND METHODS

### Plant material

Col-0 ecotype wildtype, *nrpe1/nrpd1b-11* (Pontes et al., 2006), *ago4-1* (introgressed into the Col-0 background (Wierzbicki et al., 2009)), *idn2-1* (Ausin et al., 2009), *drm2-2* (SAIL_70_E12, (Wierzbicki et al., 2008), *swi3b-2* (GABI_302G08, (Sarnowski et al., 2005)), *cmt3-11* (SALK_148381) and *met1-3* (Saze et al., 2003) were grown at 22°C under white LED light in 16h/8h day/night cycle.

### Antibodies

Rabbit polyclonal anti-histone H3 antibody (ab1791) and mouse monoclonal anti-H3K9me2 antibody (ab1220) were obtained from Abcam.

### MNase H3 ChIP-seq

2g of approximately 3.5-week old *Arabidopsis thaliana* mature leaf tissue, which was crosslinked with 0.5% formaldehyde, was ground in liquid nitrogen. MNase H3 ChIP of Col-0, *met1, cmt3* and *drm2* was carried out as described previously (Zhu et al., 2013). MNase H3 ChIP of Col-0, *nrpe1* and *swi3b* was carried out using the following protocol. Cold nuclei isolation buffer I (10 mM Tris HCl pH8, 10mM MgCl_2_, 0.4 M sucrose, 0.035% β-mercaptoethanol, 1mM phenylmethylsulfonyl fluoride (PMSF)) was added. Tissue was resuspended by vigorous vortexing and shaking. Sample was filtered using Miracloth into new 50 ml tube on ice. Miracloth was washed with 10 ml of nuclei isolation buffer I. Sample was centrifuged 15 min, 4000 g, 4°C.

Supernatant was discarded and nuclei pellet was resuspended using 1 ml of cold nuclei isolation buffer II (10 mM Tris HCl pH8, 10 mM MgCl_2_, 0.4 M sucrose, 1% Triton X-100, 0.035% β-mercaptoethanol, 1mM phenylmethylsulfonyl fluoride (PMSF), 0.02 tab/ml cOmplete EDTA-free, 0.004 mg/ml Pepstatin A). Sample was transferred to 1.5 ml tube and centrifuged for 5 min, 2000 g, 4°C. This step was repeated two more times. Pellet was resuspended using 300 μl of cold Nuclei isolation buffer II and layered on top of cold 900 ml Nuclei isolation buffer III (10 mM Tris HCl pH8, 2 mM MgCl_2_, 1.7 M sucrose, 0.15% Triton X-100, 0.035% β-mercaptoethanol, 1 mM phenylmethylsulfonyl fluoride (PMSF), 0.02 tab/ml cOmplete EDTA-free, 0.004 mg/ml Pepstatin A) in 1.5 ml tube. Sample was centrifuged for 30 min, 16000 g, 4°C and supernatant was discarded.

Isolated nuclei were washed twice with Micrococcal Nuclease (MNase) reaction buffer (10 mM Tris HCl pH8, 15 mM NaCl, 60 μM KCl, 1mM CaCl_2_) and resuspended in the same buffer. MNase enzyme (NEB; 200 Kunitz unit/μl) was added and samples were mixed by vortexing. Samples were digested for 10 minutes at 30°C. 1 volume of MNase stop buffer (30 mM Tris HCl pH8, 225 mM NaCl, 10 mM ethylenediaminetetraacetic acid (EDTA), 10 mM egtazic acid (EGTA), 0.2% sodium dodecyl sulphate (SDS), 2% Tween 20) was then added to stop the reaction. To release the chromatin from the nuclei, the sample was vortexed vigorously 5 times and centrifuged for 10 min, 14000 g. The supernatant was then transferred to a new tube. Samples for H3 ChIP were then diluted in 1 volume ChIP dilution buffer (16.7 mM Tris HCl pH8, 1.2 mM ethylenediaminetetraacetic acid (EDTA), 167 mM NaCl, 1.1% Triton X-100, 1 mM phenylmethylsulfonyl fluoride (PMSF), 0.02 tab/ml cOmplete EDTA-free, 0.004 mg/ml Pepstatin A). H3 antibody was added and sample was incubated 12-16 hours, 4°C with rotation.

Protein A magnetic beads (Pierce™) were washed three times with IP buffer (50 mM HEPES pH7.5, 150 mM NaCl, 10 μM ZnSO_4_, 1% Triton X-100, 0.05% sodium dodecyl sulphate (SDS), 1 mM phenylmethylsulfonyl fluoride (PMSF), 0.02 tab/ml cOmplete EDTA-free, 0.004 mg/ml Pepstatin A) and resuspended in 50 μl IP buffer. Beads were added to IP sample and incubated for 1 hour, 4°C with rotation. Immunoprecipitated chromatin bounded to magnetic beads was collected using magnetic separator. Beads were washed 5 min with cold buffers: two times with low salt buffer (20 mM Tris HCl pH8, 2 mM ethylenediaminetetraacetic acid (EDTA), 150 mM NaCl, 1% Triton X-100, 0.1% sodium dodecyl sulphate (SDS)), once with high salt buffer (20 mM Tris HCl pH8, 2 mM ethylenediaminetetraacetic acid (EDTA), 0.5 M NaCl, 1% Triton X-100, 0.1% sodium dodecyl sulphate (SDS)), once with LiCl buffer (20 mM Tris HCl pH8, 2 mM ethylenediaminetetraacetic acid (EDTA), 250 mM LiCl, 1% NP-100, 1% sodium deoxycholate)) and twice with TE buffer (10 mM Tris HCl pH8, 1 mM ethylenediaminetetraacetic acid (EDTA)). After the last wash, samples were transferred into new a tube and beads were collected using a magnetic separator.

For library preparation, magnetic beads were incubated with 100 μl Elution buffer (10 mM Tris HCl pH8, 1 mM ethylenediaminetetraacetic acid (EDTA), 1% sodium dodecyl sulphate (SDS)) in a thermomixer (65°C, 1400 rpm, 30 min). Beads were collected using magnetic separator and supernatant was transferred into new tube. Step was repeated and both supernatants combined. IP samples were decrosslinked by Proteinase K treatment (5 μl, 65°C, 12 h). Samples were purified using QIAquick^®^ PCR Purification Kit (35 μl of EB buffer were used). Library for Illumina sequencing was prepared using either MicroPlex Library Preparation™ Kit (Diagenode) according manufacturer instruction, using inhouse library preparation based on Bowman *et al* (Bowman et al., 2013), or prepared by the University of Michigan Advanced Genomics Core. MNase ChIP-seq experiments were performed in two biological replicates and sequenced by either 50 bp or 150 bp paired-end sequencing at the University of Michigan Advanced Genomics Core.

### MNase H3 & H3K9me2 ChIP-qPCR

Nuclei were extracted from 2g of approximately 3.5-week old *Arabidopsis thaliana* mature leaf tissue which was crosslinked with formaldehyde [0.5%] as described previously(Zhu et al., 2013) and were digested with Micrococcal Nuclease (MNase; NEB) for 10 minutes at 30°C. MNase-digested chromatin was immunoprecipitated with anti-histone H3 antibody or anti-H3K9me2 antibody. DNA was purified and used for qPCR analysis. MNase ChIP-qPCR experiments were performed in three biological replicates with region-specific primers listed in *Supplementary table 1*.

### Whole genome bisulfite-seq

Genomic DNA was isolated from approximately 3.5-week old *Arabidopsis thaliana* mature leaf tissue of Col-0 wild type, *swi3b*/+ and *nrpe1* using DNeasy Plant Mini Kit (QIAGEN). DNA was processed for bisulfite treatment and library generation at the University of Michigan Advanced Genomics Core.

### Bioinformatic data analysis

MNase H3 ChIP-seq paired-end reads from two independent biological replicates were aligned and processed to the *Arabidopsis* TAIR10 genome with Bowtie2 (Langmead and Salzberg, 2012). Mapped reads were deduplicated using PICARD tools (http://broadinstitute.github.io/picard) and filtered by fragment length between 120-170 bp and MAPQ value of >=2. Differential nucleosomes were identified using DANPOS2 (Chen et al., 2013) by filtering nucleosomes with more than 2 fold enrichment in either in Col-0 for PolV stabilized nucleosomes or in *nrpe1* for PolV destabilized nucleosomes and FDR < 0.05. Nucleosomes are then filtered using the negative binomial test with reads from biological replicates using the NBPseq R package (Di et al., 2011). For subsequent analysis we selected nucleosomes which showed more than 2 fold-change and FDR < 0.05. We further refined the nucleosome positions for well-positioned nucleosomes by filtering for main peak nucleosomes using iNPS (Chen et al., 2014). Nucleosome data was (RPM) normalized and visualized on heatmaps and profiles by calculating the number of reads using BEDTools 2.15.0 at nucleosome dyads (Quinlan and Hall, 2010). Overlap analyses with nucleosomes were performed with 1000 permutated genomic regions to obtain expected numbers and p values. SWI3B-stabilized nucleosomes were filtered for higher read counts in Col-0 than the *swi3b* mutant and an FDR<0.05. These nucleosomes were then further filtered using the negative binomial test with reads from biological duplicates using NBPseq and the nucleosomes with FDR<0.01 were selected for further analysis.

Hypermethylated nucleosomes were identified by filtering for nucleosomes with higher DNA methylation level in CHH-context inside the nucleosomes (140bps), compared to their adjacent DNA linker regions (30bps upstream and downstream of nucleosomes). The nucleosomes were filtered for the presence of more than 8 CHH-context cytosines within the nucleosomes and more than 2 CHH-context Cs in each of the adjacent linkers to correct for the sizes of the regions and frequencies of Cs. Hypomethylated nucleosomes were similarly identified, except these regions had higher levels of CHH-context DNA methylation in the adjacent DNA linker regions than the nucleosomes.

The sequencing reads from whole genome bisulfite-seq datasets were mapped to the TAIR10 genome using the Bismark software allowing no mismatches (Krueger and Andrews, 2011). DNA methylation levels were calculated by the ratio of #C/(#C+#T) after selecting for Cs with at least 5 sequenced reads. Differentially Methylated Regions (DMRs) were identified using methylKit package in R (Akalin et al., 2012). The bin sizes used were 100bp bins with a step-size of 50bps. 10 minimum bases were required in each tile. A 10% minimum methylation difference was selected for in each of the tiles and an FDR value of 0.01 was used. The number of MNase-H3 ChIP-seq reads overlapping these DMRs were then plotted as a profile.

### Other datasets used in this study

*Arabidopsis* genome annotations (TAIR10) were obtained from TAIR (www.arabidopsis.org). Pol V ChIP-seq data (SRA054962) and peak list and Pol V RIP-seq data (GSE70290) and annotated regions were published previously (Böhmdorfer et al., 2016; Wierzbicki et al., 2012). DNA methylation data from *idn2*, *ago4*, *drm2*, *met1* and *cmt3* mutants as well as corresponding Col-0 and *nrpe1* controls were obtained from GSE39901 (Stroud et al., 2013).

### Data deposition

High throughput sequencing datasets obtained in this study have been deposited in Gene Expression Omnibus under accession GSE148173.

## ACKNOWLEDGMENTS

Research reported in this publication was supported by the National Institute of General Medical Sciences of the National Institutes of Health under award number R01GM108722 to ATW. JD was supported by the National Science Center grants Etiuda UMO-2014/12/T/NZ2/00246 and Opus UMO-2017/25/B/NZ1/00603.

## SUPPLEMENTAL FIGURE LEGENDS

**Table S1:**
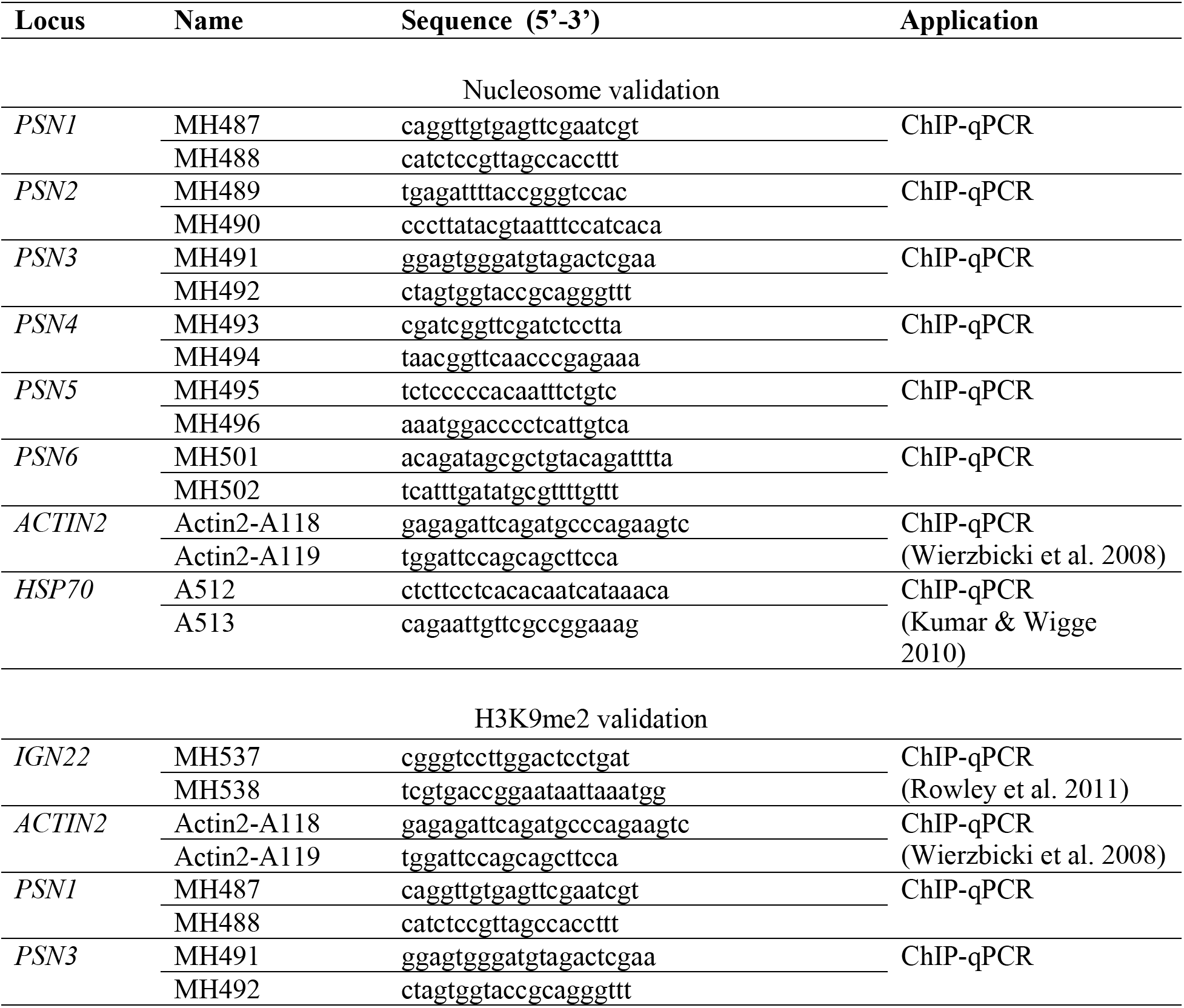
Oligonucleotides used in this study.

**Figure S1:**
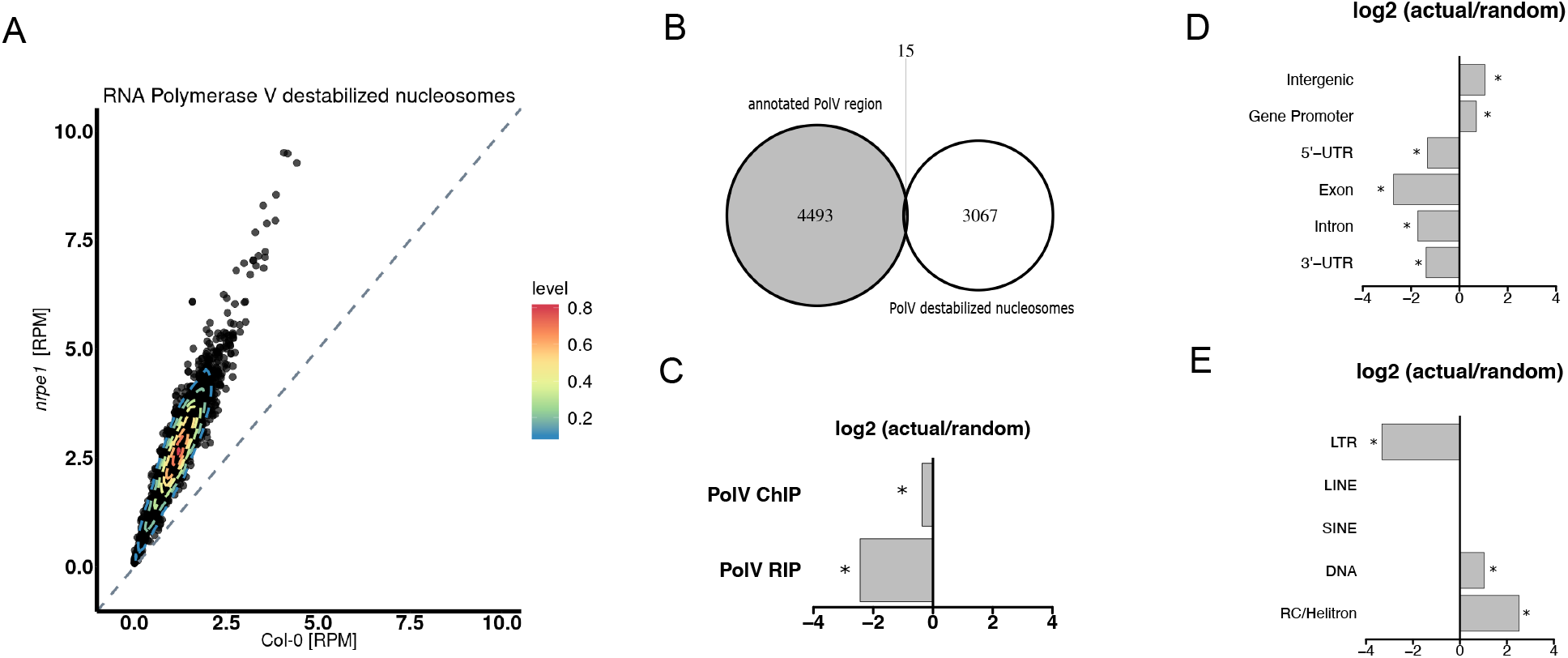
Pol V affects nucleosomes by a combination of direct and indirect mechanisms. A. MNase H3-ChIP seq signal on Pol V destabilized nucleosomes. B. Overlap between Pol V destabilized nucleosomes and annotated Pol V transcribed regions. C. Enrichment of Pol V destabilized nucleosomes on annotated Pol V transcribed or bound regions (random permutation test; 1000 iterations; p-value < 0.001). D. Enrichment of Pol V stabilized nucleosomes on various genomic regions (random permutation test; 1000 iterations; p-value < 0.001). E. Enrichment of Pol V stabilized nucleosomes on annotated transposable element regions (random permutation test; 1000 iterations; p-value < 0.001).

**Figure S2:**
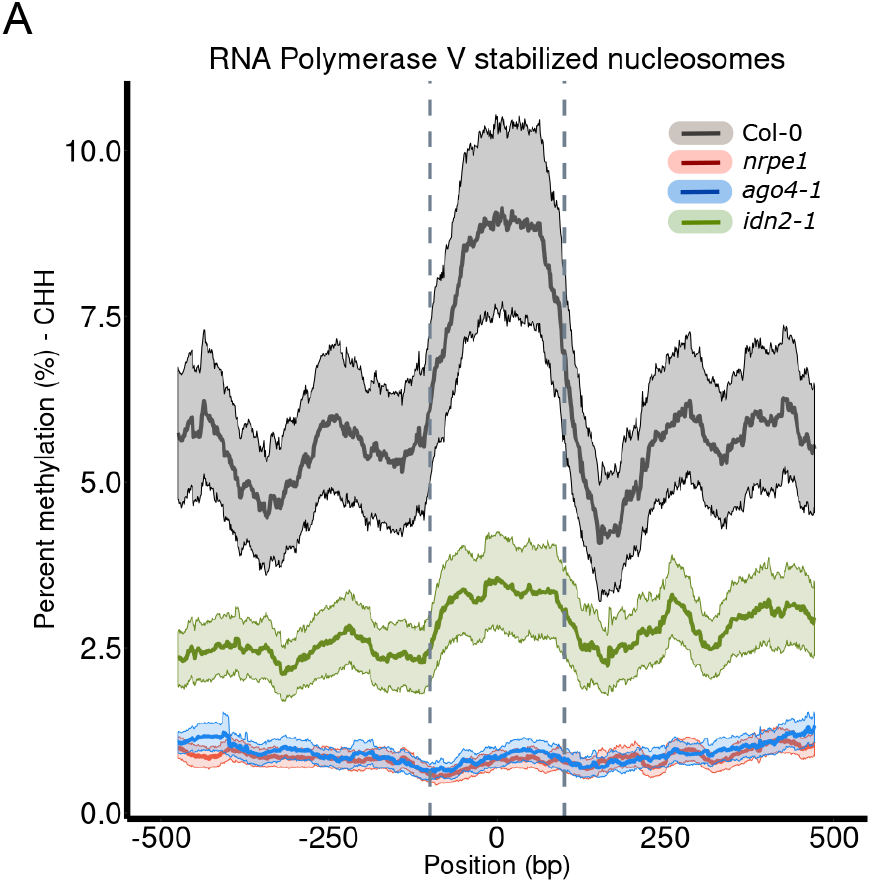
IDN2 connects siRNA and lncRNA to nucleosome positioning. A. Average levels of CHH methylation on and around Pol V stabilized nucleosomes dyads using datasets from Stroud et al (2013).

**Figure S3:**
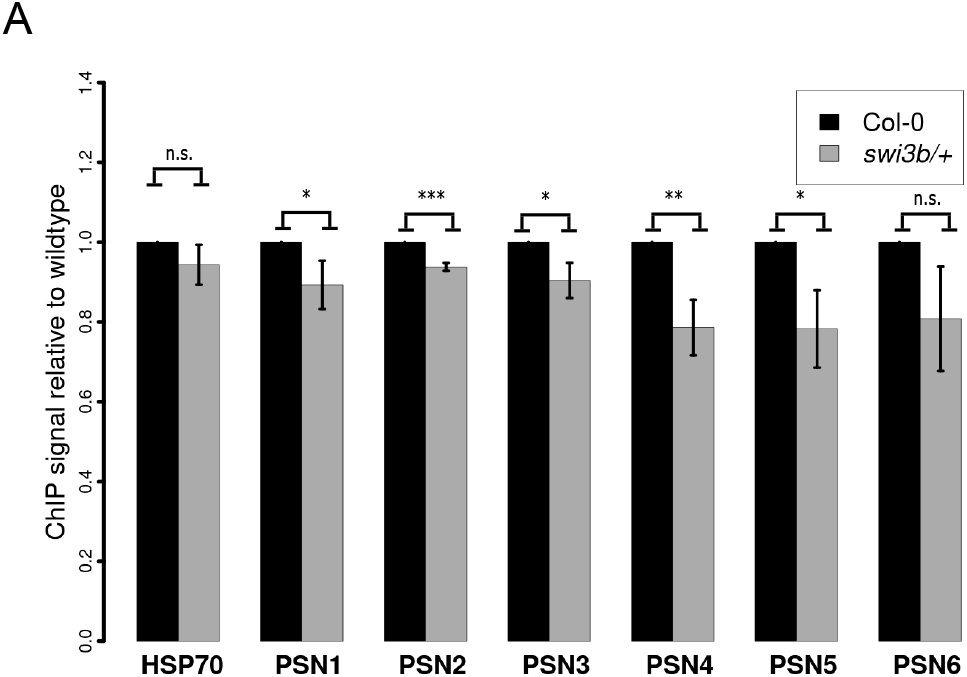
Pol V positions nucleosomes through the SWI/SNF complex. A. Locus-specific validation of Pol V stabilized nucleosomes by MNase H3 ChIP followed by qPCR. Significance tested using t-test (n.s. = not significant, * = p-value < 0.05, ** = p-value < 0.01,*** = p-value < 0.001). ChIP signal values were normalized to ACTIN2 and Col-0 wildtype. Error bars indicate standard deviations from three biological replicates.

**Figure S4:**
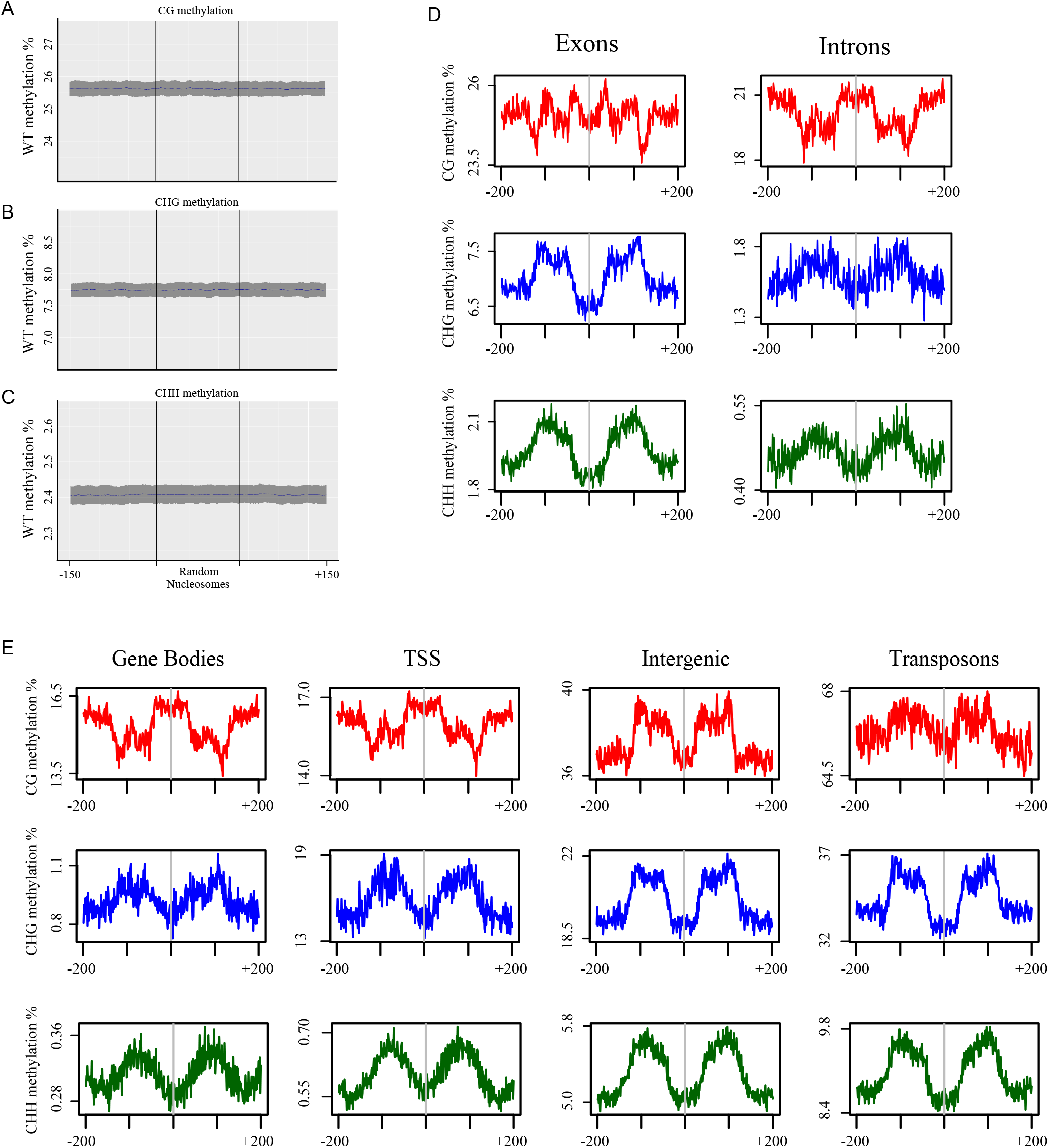
Preferential methylation of linker DNA. A. Average levels of CG methylation at random nucleosome-sized regions. Ribbon indicates confidence intervals with p < 0.05. B. Average levels of CHG methylation at random nucleosome-sized regions. Ribbon indicates confidence intervals with p < 0.05. C. Average levels of CHH methylation at random nucleosome-sized regions. Ribbon indicates confidence intervals with p < 0.05. D. Average levels of CG, CHG and CHH methylation at and around annotated nucleosomes overlapping exons or introns. E. Average levels of CG, CHG and CHH methylation at and around annotated nucleosomes overlapping gene bodies, TSS, intergenic regions and transposable elements.

**Figure S5:**
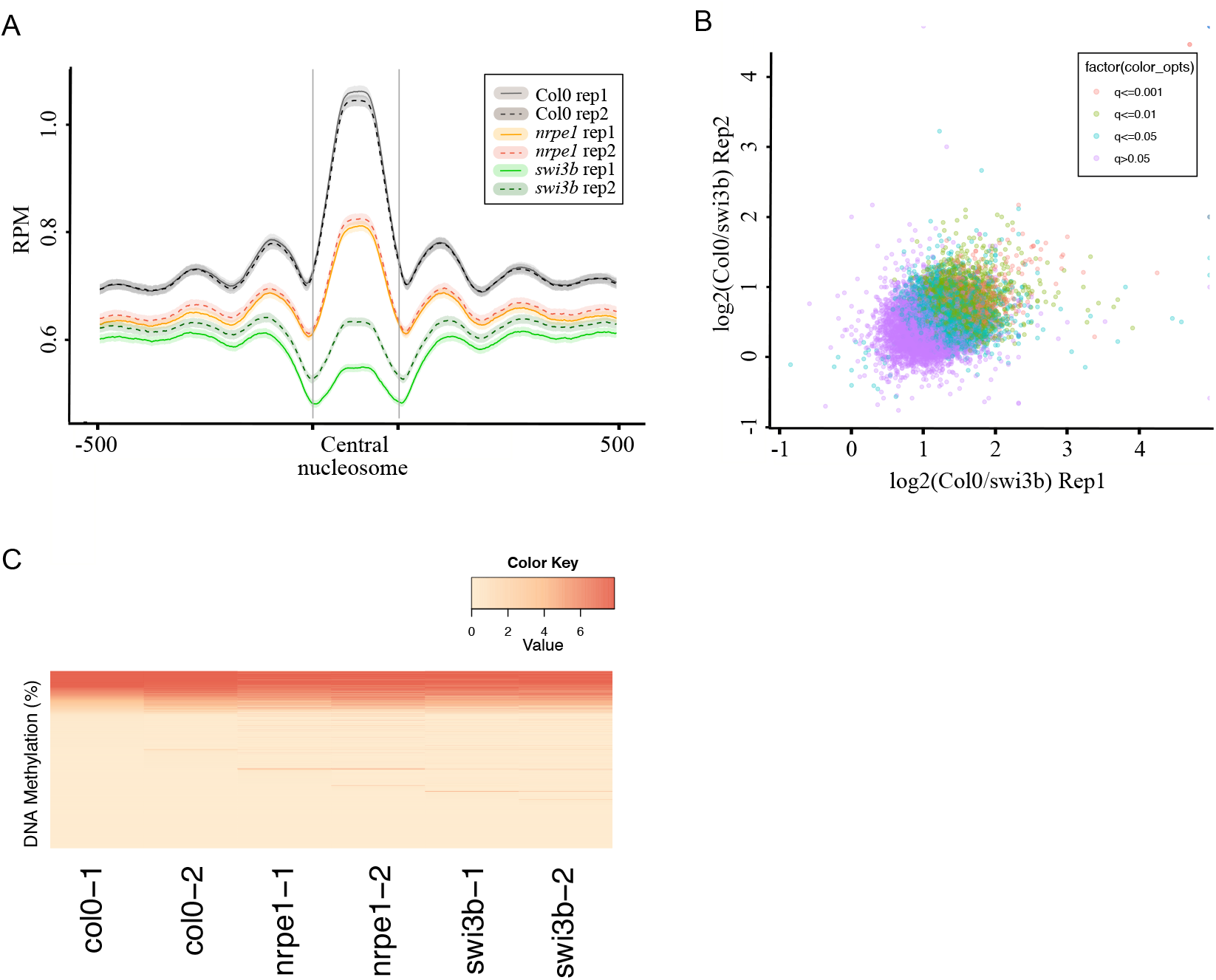
SWI/SNF complex is not required for DNA methylation on positioned nucleosomes. A. Average levels of MNase H3 ChIP in two biological replicates of Col-0, *nrpe1* and *swi3b*/+ at and around SWI3B stabilized nucleosomes. B. Comparison of biological replicates of MNase H3 ChIP in Col-0 and *swi3b*/+. Colors scale represents FDR cutoff values. C. DNA methylation levels at individual SWI3B stabilized nucleosomes.

**Figure S6:**
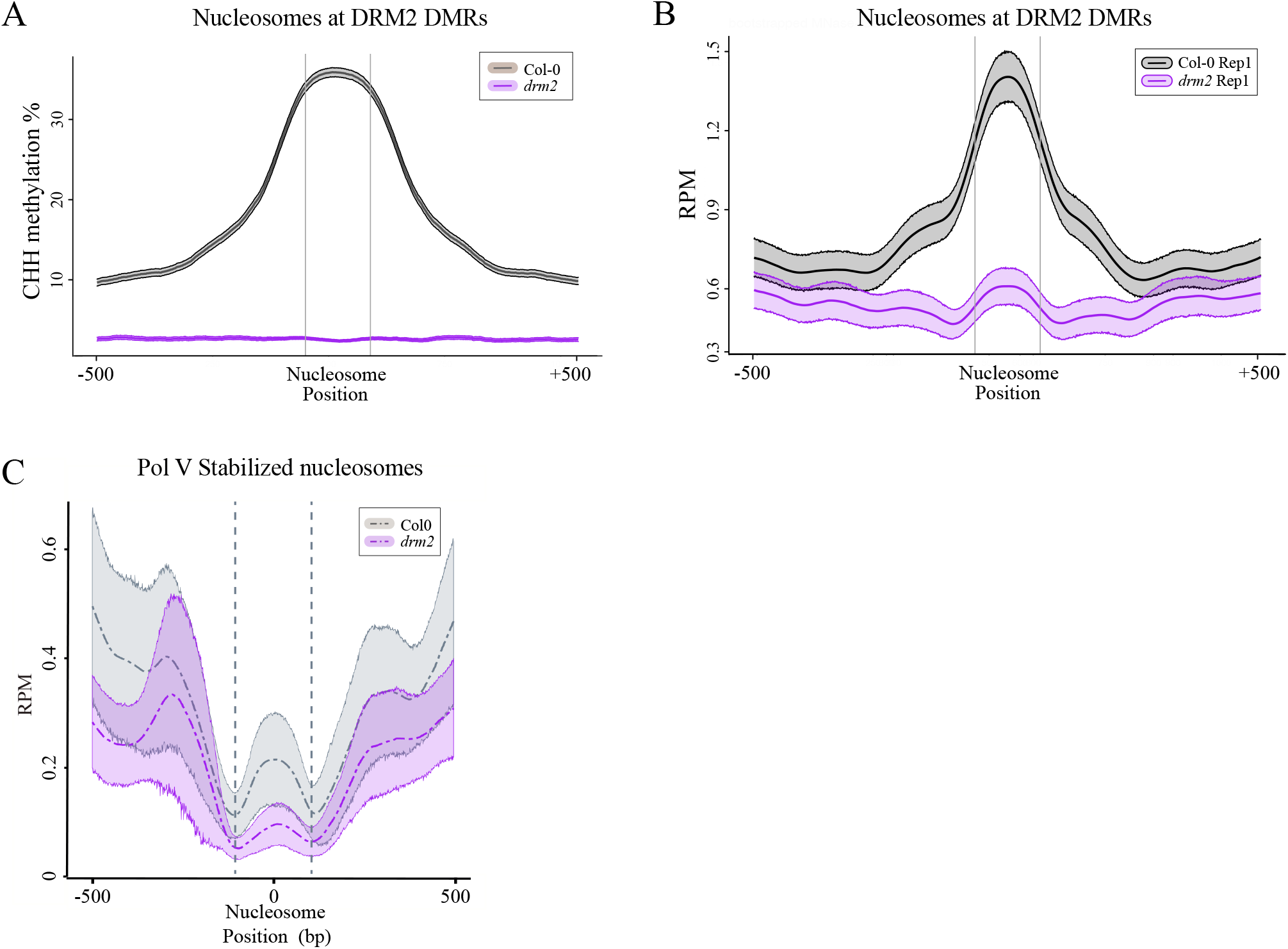
DNA methylation is needed for positioning nucleosomes at differentially methylated regions. A. Average levels of CHH methylation at and around annotated nucleosomes that overlap DRM2 DMRs. Ribbons indicate confidence intervals with p < 0.05. B. Average levels of MNase H3 ChIP signal at nucleosomes overlapping DRM2 DMRs. Ribbons indicate confidence intervals with p < 0.05. C. Average levels of MNase H3 ChIP signal at Pol V stabilized nucleosomes. Ribbons indicate confidence intervals with p < 0.05.

